# Mechanical force regulates ligand binding and function of PD-1

**DOI:** 10.1101/2023.08.13.553152

**Authors:** Kaitao Li, Paul Cardenas-Lizana, Anna V. Kellner, Zhou Yuan, Eunseon Ahn, Jintian Lyu, Zhenhai Li, Khalid Salaita, Rafi Ahmed, Cheng Zhu

**Affiliations:** Wallace H. Coulter Department of Biomedical Engineering; Parker H. Petit Institute for Bioengineering and Biosciences; George W. Woodruff School of Mechanical Engineering, Georgia Institute of Technology, Atlanta, Georgia 30332, USA; Emory Vaccine Center; Department of Microbiology and Immunology, Emory University School of Medicine; Department of Chemistry, Emory University, Atlanta, GA 30322

## Abstract

Immune checkpoint blockade targeting PD-1 shows great success in cancer therapy. However, the mechanism of how ligand binding initiates PD-1 signaling remains unclear. As prognosis markers of multiple cancers, soluble PD-L1 is found in patient sera and can bind PD-1, but fails to suppress T cell function. This and our previous observations that T cells exert endogenous forces on PD-1– PD-L2 bonds prompt the hypothesis that mechanical force might be critical to PD-1 triggering, which is missing in the soluble ligand case due to the lack of mechanical support afforded by surface-anchored ligand. Here we show that PD-1 function is eliminated or reduced when mechanical support on ligand is removed or dampened, respectively. Force spectroscopic analysis reveals that PD-1 forms catch bonds with both PD-Ligands <7 pN where force prolongs bond lifetime, but slip bonds >8 pN where force accelerates dissociation. Steered molecular dynamics finds PD-1–PD-L2 complex very sensitive to force due to the two molecules’ “side-to-side” binding via β sheets. Pulling causes relative rotation and translation between the two molecules by stretching and aligning the complex along the force direction, yielding new atomic contacts not observed in the crystal structure. Compared to wild-type, PD-1 mutants targeting the force-induced new interactions maintain the same binding affinity but display lower rupture force, shorter bond lifetime, reduced tension, and most importantly, impaired capacity to suppress T cell activation. Our results uncover a mechanism for cells to probe the mechanical support of PD-1–PD-Ligand bonds using endogenous forces to regulate PD-1 triggering.

## Main

Programmed cell death-1 (PD-1) is an immune checkpoint receptor and a hallmark of T cell exhaustion in chronic viral infection and cancer^1-3^. PD-1 consists of a single IgV domain, a ∼20 AA stalk, a transmembrane region and an intracellular tail with two tyrosine-based signaling motifs: an immunoreceptor tyrosine-based inhibitory motif (ITIM) and an immunoreceptor tyrosine-based switch motif (ITSM)^4^. Binding of PD-1 to its ligand (PD-L1 or PD-L2) triggers the phosphorylation of ITIM and ITSM, which recruits and activates SH2-containing tyrosine phosphatase 2 (SHP-2). Activated SHP-2 dephosphorylates a panel of signaling molecules downstream of TCR and CD28, thereby suppressing T cell activation and function^5,6^. Blocking antibodies targeting PD-1–PD-L1 interaction have yielded great success in cancer immunotherapy^7,8^. However, for a molecule like PD-1 with such a simple structure, the detailed molecular mechanism as how it transduces ligand binding to initiate signaling remains unclear. It becomes even more puzzling that as a prognosis marker of various types of cancer, some soluble PD-L1 splicing variants retain their ability of PD-1 binding but fail to trigger suppression of T cells^9-12^. We also found in this study that soluble PD-Ligands, even in tetrameric forms, cannot induce PD-1 function whereas cell surface or bead-coated PD-Ligands delivered robust PD-1 triggering.

It is worth noting that one of the critical components missing in the soluble form comparing with the surface-anchored form of ligands is the ability to provide physical support to the engaged PD-1–PD-Ligand bond for it to bear mechanical load. Using DNA-based molecular tension probes (MTP) with a locking strand to accumulate the force signals we have shown that activated T cells actively apply forces on PD-1 engaged with PD-L2 or anti-PD-1 antibody^13^. In this study we also observed significant tension using MTP of 4.7 pN threshold force presenting PD-L1 or PD-L2. The formation and movement of PD-1 microclusters^5,14^ as well as the ability of PD-1–PD-L2 interaction to drive “synapse”-like interface between CHO cells^15^ also suggest PD-1–PD-Ligand bonds are constantly under mechanical load. Emerging evidence shows that mechanical force plays a critical role in immune cell functions by modulating the binding and signaling of various immune receptors^16-19^. For example, force can enhance antigen recognition by TCR and BCR and potentiate target killing by effector T cells^20-25^. At the molecular level, TCR antigen recognition is enhanced by the dynamic responses of TCR–pMHC interaction to force application – potent ligands form catch bonds with more durable TCR engagement and signaling, whereas weak ligands form slip bonds that readily rapture under force^20,22,26-29^. Together, these findings raise the intriguing question of the role of force on PD-1 ligand binding and function.

In this study, we analyzed the effect of force on PD-1–PD-Ligand bonds combining *in situ* kinetic measurements using biomembrane force probe (BFP) and *in silico* simulations using steered molecular dynamics (SMD). We found that force elicits catch-slip bonds for both PD-1–PD-L1 and PD-1–PD-L2 interactions, where forces below 7 pN prolongs bond lifetime (catch) and forces above 8 pN accelerates dissociation (slip). Corroborating the force-enhanced bonding, SMD reveals that pulling on the two C-termini of the PD-1–PD-L2 complex induces large relative rotation and translation between the two molecules, which is accompanied by formation of new atomic contacts not observed in the crystal structure in absence of applied force. Mutating residuals on PD-1 involved in the force-induced new non-covalent interactions decreased the mechanical stability of the PD-1–PD-L2 bond manifesting a lower force required for bond rupture, shorter bond lifetime under sustained force, and reduced the number of bonds bearing above threshold forces, despite the lack of effect of the mutations on the *in situ* PD-1–PD-L2 binding affinity in the absence of force. Most importantly, these PD-1 mutants demonstrate impaired ability to suppress TCR-CD3 induced NFκB activation. Overall, our results suggest force critically regulates the mechanical stability of PD-1–PD-Ligand bond, which is essential for efficient PD-1 triggering.

### The inhibitory function of PD-1 requires ligand bearing mechanical support

Recent studies have found soluble forms of alternatively spliced PD-1 ligands as prognosis biomarkers in various cancers^10,30^. Despite their ability to bind to PD-1, results were discrepant regarding whether soluble PD-1 ligands can induce immunosuppression^9,11,12,31^. To resolve this, we compared PD-1 function triggered by PD-1 ligands expressed on cell membrane or immobilized on beads versus in soluble tetrameric forms. NFκB::eGFP reporter Jurkat cells^32^ were transduced to overexpress a chimeric PD-1 molecule consisting of mouse PD-1 ectodomain (Met1 to Met169) and human PD-1 transmembrane and intracellular domain (Val171 to Leu288) (Fig. S1A). GFP expression can be induced by stimulating the cells with anti-CD3 (clone OKT3) full antibody or T-cell stimulator cells (TSC) that expresses a membrane-anchored single-chain fragment variable (scFv) of OKT3^33^. Plain and PD-1 reporter Jurkat cells were cocultured with control or PD-Ligand expressing TSCs (Figs. S1B&C) and analyzed for their GFP expression (Figs. 1A-B). As expected, cell membrane PD-L1 or PD-L2 triggered potent inhibition of OKT3-induced GFP expression as quantified by the frequency of GFP^+^ PD-1 reporter cells and their geometric mean fluorescence intensity (Figs. 1B-D). To avoid altering anti-CD3 presentation when comparing immobilized *vs* soluble PD-Ligands, plain or PD-1 reporter cells were stimulated with soluble OKT3 together with soluble or bead-coated PD-Ligands and analyzed for their GFP expression (Figs. 1E-F). Comparing to control group of anti-CD3 plus soluble streptavidin (SA), the experimental groups of anti-CD3 plus PD-L1 or PD-L2 tetramer failed to induce suppression of GFP induction in PD-1 reporter cells just like in plain control cells, despite the high concentration (20 μg/ml based on monomer) used (Figs. 1G-H). However, when coated on beads, both PD-L1 and PD-L2 triggered potent inhibition of anti-CD3 induced GFP expression in PD-1 reporter cells but not in plain control cells (Figs. 1G-H). Together, these results indicated that successful triggering of PD-1 requires cell membrane or surface anchored ligands, regardless of their ability to cluster by crosslinking. Interestingly, such a requirement is not limited to natural ligands of PD-1. When PD-Ligands in Figs. 1E&F were replaced with anti-PD-1 antibodies, all three clones (29F.1A12, J43, and RMP1-30) show no effect of PD-1 triggering in soluble tetramer form but strong effect on PD-1 triggering when immobilized on beads (Fig. S2), suggesting surface anchoring is essential to the PD-1 agonist effect.

**Fig. 1.**
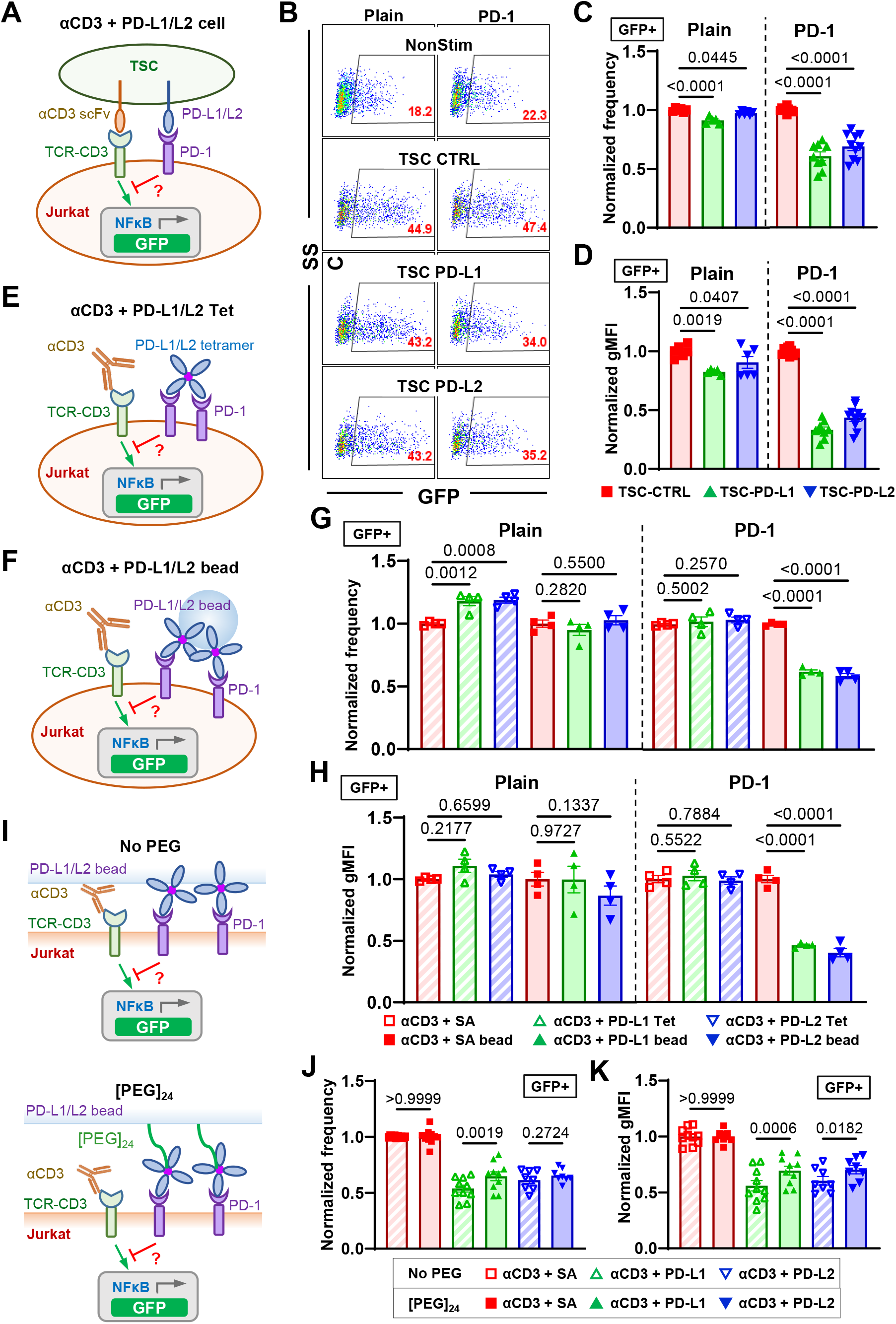
The inhibitory function of PD-1 requires PD-Ligands to bear mechanical support. (A) Schematics of stimulating NFκB::eGFP reporter Jurkat cells with T-cell stimulator cells (TSC) expressing a single-chain fragment variable (scFv) of anti-CD3 (clone OKT3) and PD-L1 or PD-L2. (B) Representative SSC *vs* GFP plots of reporter Jurkat cells 24 hr after stimulation with indicated conditions. (C-D) Quantification of GFP expression for conditions in B. n = 5 - 6 for plain pooled from 3 independent experiments or n = 9 - 10 for PD-1 reporter cells pooled from 5 independent experiments. (E-F) Schematics of stimulating NFκB::eGFP reporter Jurkat cells with soluble anti-CD3 and soluble (E) or bead-coated (F) PD-Ligands. (G-H) Quantification of GFP expression for conditions in E&F. n = 4 for all conditions pooled from two independent experiments. (I) Schematics of stimulating NFκB::eGFP reporter Jurkat cells with soluble anti-CD3 and PD-Ligands coated beads without or with [PEG]_24_ spacer arm. (J-K) Quantification of GFP expression for conditions in I. n = 10, 10, and 8 for SA, PD-L1, and PD-L2, respectively, pooled from 5 independent experiments. Normalized frequency (C, G, and J) and normalized geometric mean fluorescence intensity (gMFI) (D, H, and K) were calculated as (sample – averaged background)/(anti-CD3 control – averaged background). Numbers on graphs represent p values calculated from two-tailed student t-test.

Given that force has been shown to have critical effect in the triggering of other membrane receptors^16-19^, the above data prompt us to hypothesize that one critical component for PD-1 triggering is mechanical force on PD-1, which can be generated endogenously by T cells and supported by engaged ligands or antibodies if they are anchored on a solid surface to provide counter-balance force^13^. To further test this hypothesis, we altered the bead-based ligand presentation system to modulate its ability to support mechanical forces on PD-L1/L2. By introducing a [PEG]_24_ spacer between the bead surface and SA, a ∼10 nm extra length was added to the ligand, which is comparable to the dimension of a PD-1–PD-Ligand spanned across the intercellular junction^15^. This should prevent at least part of the PD-1–PD-Ligand bonds to be fully stretched due to the ligand elongation, but should not affect PD-1–PD-Ligand binding (Fig. 1I). When both bead-coated PD-Ligands were tested against PD-1 reporter Jurkat cells under anti-CD3 stimulation, PD-Ligands with a [PEG]_24_ spacer induced less inhibition comparing to those without the spacer (Figs. 1J-K), supporting our hypothesis because PD-1 triggering was reduced by dampening the mechanical support on PD-Ligands.

### Molecular Tension probes reveal active cellular forces applied to PD-1–PD-Ligand bonds

We next used MTP-tagged PD-L1 or PD-L2 to directly measure forces applied to single PD-1– PD-ligand bonds by CHO cells overexpressing PD-1 (Fig. 2A). The MTP consists of three DNA strands linking a Cy3B fluorophore to the PD-Ligand and a BHQ2 quencher to the glass surface, which are brought together by a hairpin structure to controls the force threshold by varying length and GC content^34,35^. Forces on PD-1–PD-Ligand that are above the threshold unfold the hairpin, thereby separating the Cy3B from the BHQ2 to enable fluorescence (Fig. 2A). Reflection interference contrast microscopy (RICM) show that CHO cells spread similarly on 4.7 and 12 pN MTPs conjugated with both PD-Ligands, but epi-fluorescence imaging reveal significantly higher Cy3B signals from the 4.7 than the 12 pN MTPs (Fig. S3 and Video S1), yet both cell spreading and tension were nearly eliminated by an anti-PD-1 blocking antibody (Fig. 2B-D). These results indicate that CHO cell spreading and pulling are mediated by specific PD-1–PD-Ligand interaction. While the spreading areas are similar for PD-L1 *vs* PD-L2 or MTPs with 4.7 pN *vs* 12 pN threshold forces (Fig. 2B&C), the tension signals were stronger for PD-L2 than PD-L1 with 4.7 pN MTP but the difference vanished as the tension signals for both PD-Ligands decreased significantly when the MTP’s force threshold was increased to 12 pN (Fig. 2B&D). These data extend previous results that activated OT-1 T cells pull on PD-1 engaged PD-L2 or anti-PD-1 antibody detected using a locking strand enhanced MTP to accumulative the weak tension signals^13^. The endogenous forces exerted on PD-1 may result from PD-1’s intracellular coupling to cytoskeleton, as ligand-bound PD-1 forms microclusters and moves centripetally on T cell membrane^5,14^, and expressing PD-1 and PD-L2 in CHO cells can drive the formation of synapse-like interface with concurrent accumulation of PD-1 and PD-L2^15^. The finding that PD-1 forces are observed in CHO cells in addition to T cells suggests that such forces may be intrinsic to CHO cells irrespective of PD-1’s inhibitory function. This supports the hypothesis that PD-1–PD-Ligand bonds can sustain forces between 4.7 pN and 12 pN and that PD-1–PD-L2 bonds are more mechanically stable than PD-1–PD-L1 bonds since their affinities are similar^36,37^.

**Fig. 2.**
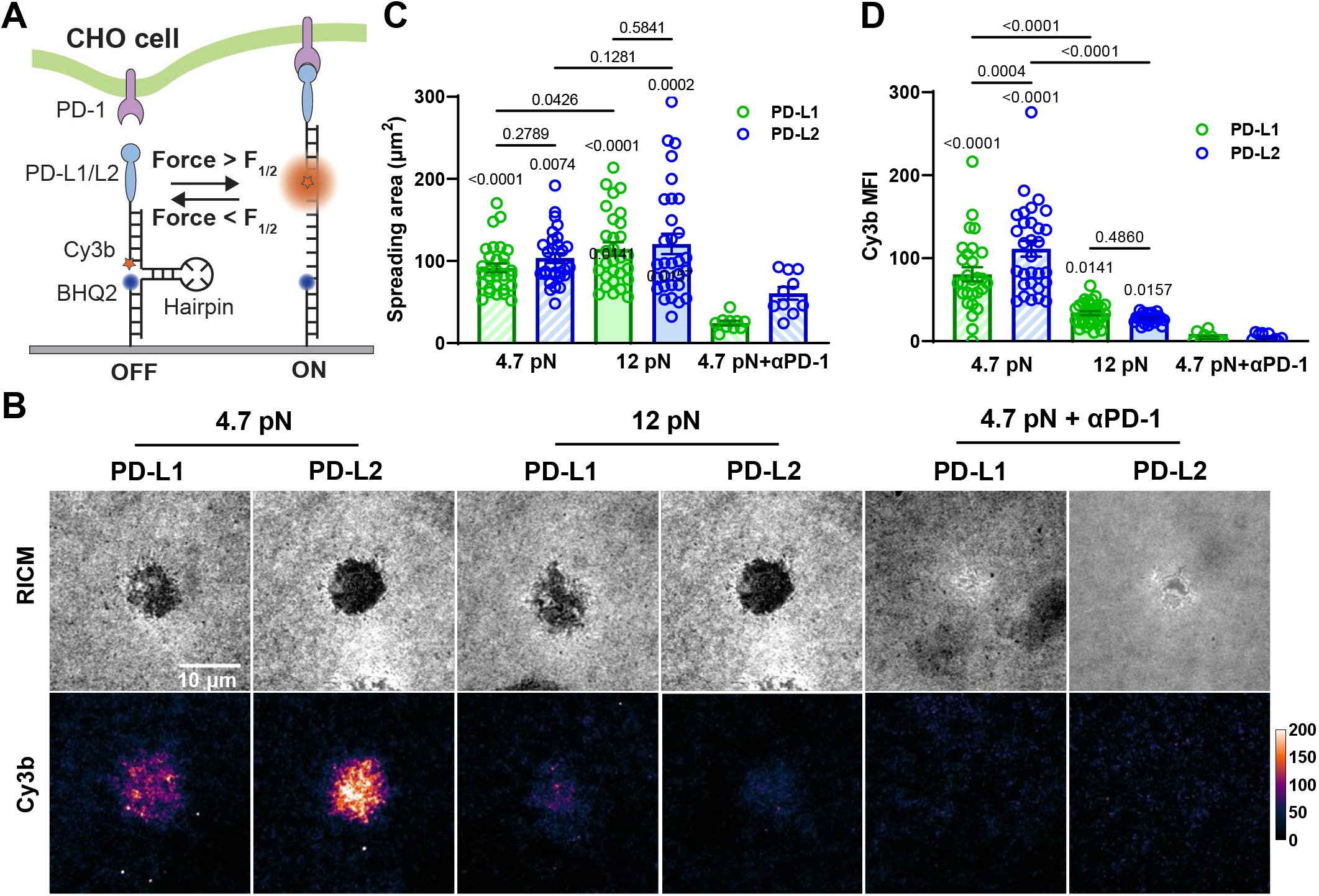
DNA-based MTPs reveal active cellular forces applied to PD-1–PD-Ligand bonds. (A) Schematics of detecting cellular forces on PD-1–PD-Ligand bonds using DNA-based molecular tension probes (MTPs). Force above unfolding threshold of the hairpin separates Cy3B from the BHQ2 quencher enabling fluorescence. (B) Representative reflection interference contrast microscopy (RICM) and Cy3b fluorescence images of PD-1 expressing CHO cells 30 min after landing on glass surface functionalized with MTPs of indicated conditions. For PD-1 blockade, cells were pre-incubated with PD-1 blocking antibody clone 29F.1A12 before imaging. (C-D) Quantification of cell spreading area (C) and tension signal (D) for conditions in B. n = 30, 30, 30, 30, 10, and 10 pooled from 3 independent experiments. Numbers on graphs represent p values calculated from two-tailed student t-test comparing two groups as indicated or each group to corresponding PD-1 blockade control.

### PD-1 forms catch bond with PD-L1 and PD-L2

To test the aforementioned hypothesis, we performed dynamic force spectroscopic analysis of the rupture forces (force-ramp spectroscopy) and bond lifetimes (force-clamp spectroscopy) of PD-1 expressed on CHO cells interacting with PD-L1 or PD-L2 using a biomembrane force probe (BFP) (Fig. 3A)^38^. A CHO cell was repetitively brought into contact with a BFP bead coated with PD-Ligand and then separated until the bond ruptured (Fig. 3B) or held at a preset force until spontaneous dissociation (Fig. 3C). The magnitude of force is calibrated from the spring constant (∼0.3 pN/nm) of the BFP and the displacement of the bead, which was tracked with millisecond temporal resolution and sub-nanometer spatial resolution, resulting in pico-newton force resolution^39^. Hundreds of bonding events were analyzed for PD-1 interacting with each PD-Ligand and were pooled to analyze the rupture force distribution at a loading rate of 1000 pN/s (Fig. 3D) or bond lifetime distribution at a force of 7 pN (Fig. 3F). The cumulative frequency of ruptured events *vs* force at which bond ruptures follows a sigmodal shape with the PD-1–PD-L2 curve right-shifted towards higher forces relative to the PD-1–PD-L1 curve (Fig. 3E). The force at which 50% of the bonds rupture (*F*_1/2_) was 12.1 pN for PD-1–PD-L2 bonds, significantly higher than the 10.3 pN value for the PD-1–PD-L1 bonds.

**Fig. 3.**
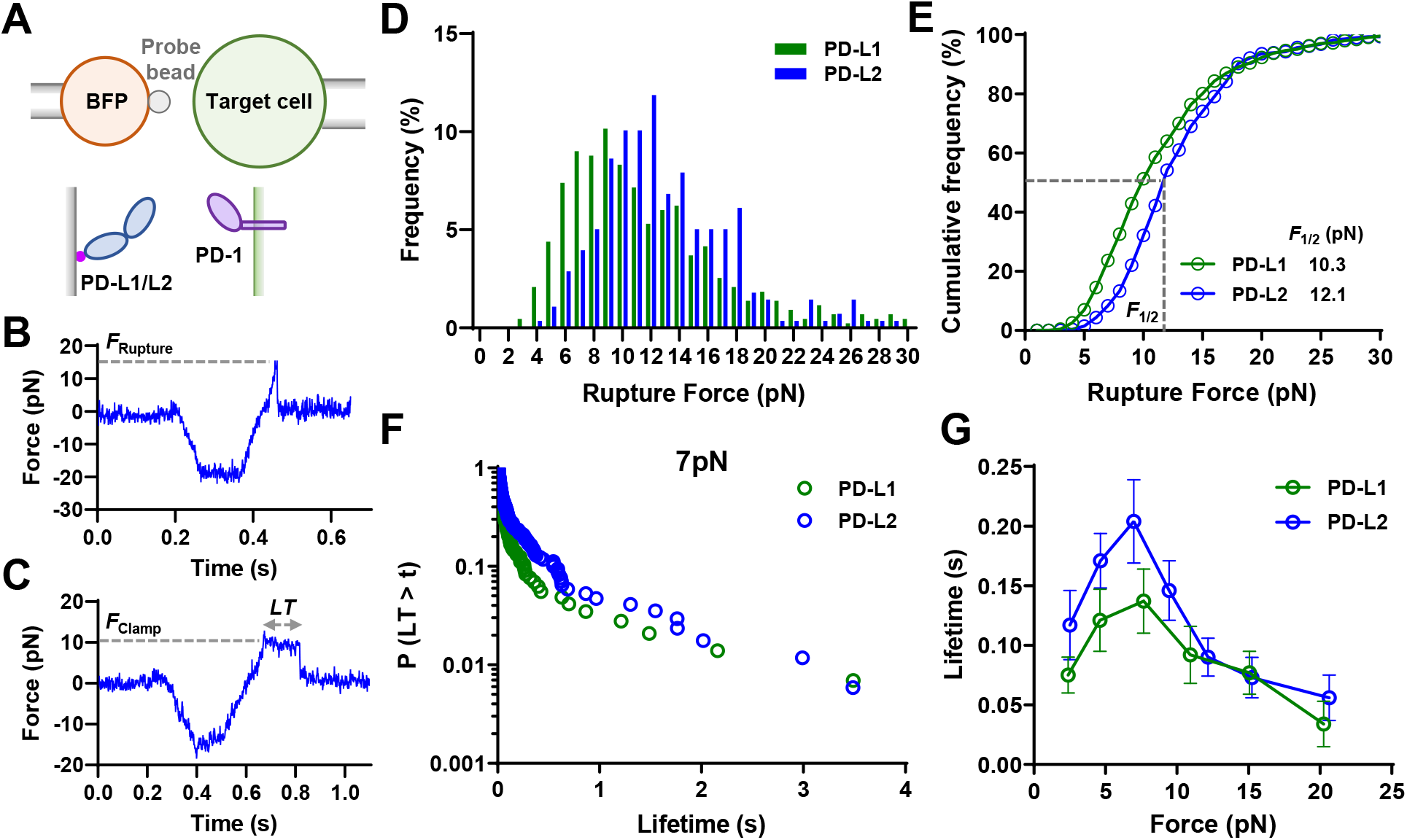
PD-1 forms catch bond with PD-L1 and PD-L2. (A) Schematics of force spectroscopic analysis of PD-1–PD-Ligand bonds using biomembrane force probe (BFP). CHO cells expressing PD-1 were analyzed against BFP bead coated with PD-L1 or PD-L2. Bead displacements were tracked with high spatiotemporal resolution and translated into force after multiplying by the spring constant of BFP. (B&C) Representative raw traces of rupture force (B) and bond lifetime (C) measurements. A target cell held by piezo-driven micropipette was brought into brief contact with a bead (approach and contact) to allow for bond formation. Upon separation the target cell either kept retracting to rupture the bond (B) or stopped and held at a predefined force level until bond dissociated spontaneously (C). (D&E) Force histograms (D) and cumulative frequencies (E) of rupture events of 433 PD-1–PD-L1 and 278 PD-1–PD-L2 bonds. *F*_1/2_ is defined as the force level at which 50% of the bonds are ruptured. p < 0.0001 comparing *F*_1/2_ of PD-1–PD-L1 and PD-1–PD-L2 using two-tailed Mann-Whitney test. (F&G) Survival frequencies at the 7 pN force bin (F) and mean ± sem bond lifetime vs force plots (G) of 497 PD-1–PD-L1 and 777 PD-1–PD-L2 bonds. p < 0.0001 comparing lifetime vs force distributions of PD-1–PD-L1 and PD-1–PD-L2 using two-tailed two-dimensional Kolmogorov-Smirnov test.

Interestingly, the mean ± sem lifetime *vs* force curves of PD-1–PD-L1 and PD-1–PD-L2 interactions both display a catch-slip bond characterstics, where bond lifetime increased with force below 7 pN (“catch”) and decreased with force above 8 pN (“slip”) (Fig. 3G), a phenomenon observed for many other molecular interactions^19,40^. Plotting the pooled bond lifetime distribution of each force bin reveals a dynamic composition of short (<0.1 s), intermediate (0.1 s to 1 s), and long (> 1s) lifetime species in response to force, suggesting multiple states during force-induced dissociation (Fig. S4&S5). Consistent with the higher rupture forces of the PD-1–PD-L2 bond than the PD-1–PD-L1 bond, the lifetime *vs* force curve for the PD-1–PD-L2 bond was up-shifted toward longer lifetime across different force levels relative to the curve for the PD-1–PD-L1 bond (Fig. 3G). These results support our hypothesis and suggest that PD-1–PD-L2 bond is more mechanically stable than PD-1–PD-L1 bond, which is consistent with the higher tension signal mediated by PD-1–PD-L2 interaction than PD-1–PD-L1 interaction (Fig. 2D).

### SMD reveals force-induced conformational changes in PD-1–PD-L2 complex and formation of new non-covalent contact at the atomic level

The structures of PD-1–PD-Ligand complexes of human and mouse species all show a “side-to-side” interaction of β-sheets from two immunoglobular domains, manifesting an assembly similar to the variable domains of α and β chains of TCR or heavy and light chains of antibody^15,41,42^. In such assembly positions, PD-1 and PD-L2 form with a sharp angle between their respective axes, providing a lever arm for the tensile force to exert a moment to unbend this angle. To gain structural insights of the effect of force on PD-1–PD-Ligand complex, we applied free molecular dynamics (FMD) and steered molecular dynamics (SMD) to simulate the dynamic responses of the PD-1– PD-L2 structure (PDB: 3BP5) without force or with force applied to the C-termini of the two molecules, respectively. Atomic coordinates were tracked for a total of 60 ns and analyzed for overall structural changes and molecular bonding events (Fig. 4). Snapshots of the complex at different time points show that force indeed gradually aligned the two molecules along their long axes converting the “side-to-side” interaction to a nearly “head-to-head” position before bond rupture (Fig. 4A and Video S2). This conformational change manifests large changes in both the relative angle (from ∼50° to ∼160°) and displacement (∼30 Ǻ measured by RMSD) of the two molecules with distinct transitions among multiple phases (Fig. 4A&B). Comparing to the direct dissociation from “side-to-side” binding under force-free condition, such force-induced multi-phase dissociation may explain the dynamic composition of short, intermediate, and long bond lifetimes under force (Fig. S3&S4).

**Fig. 4.**
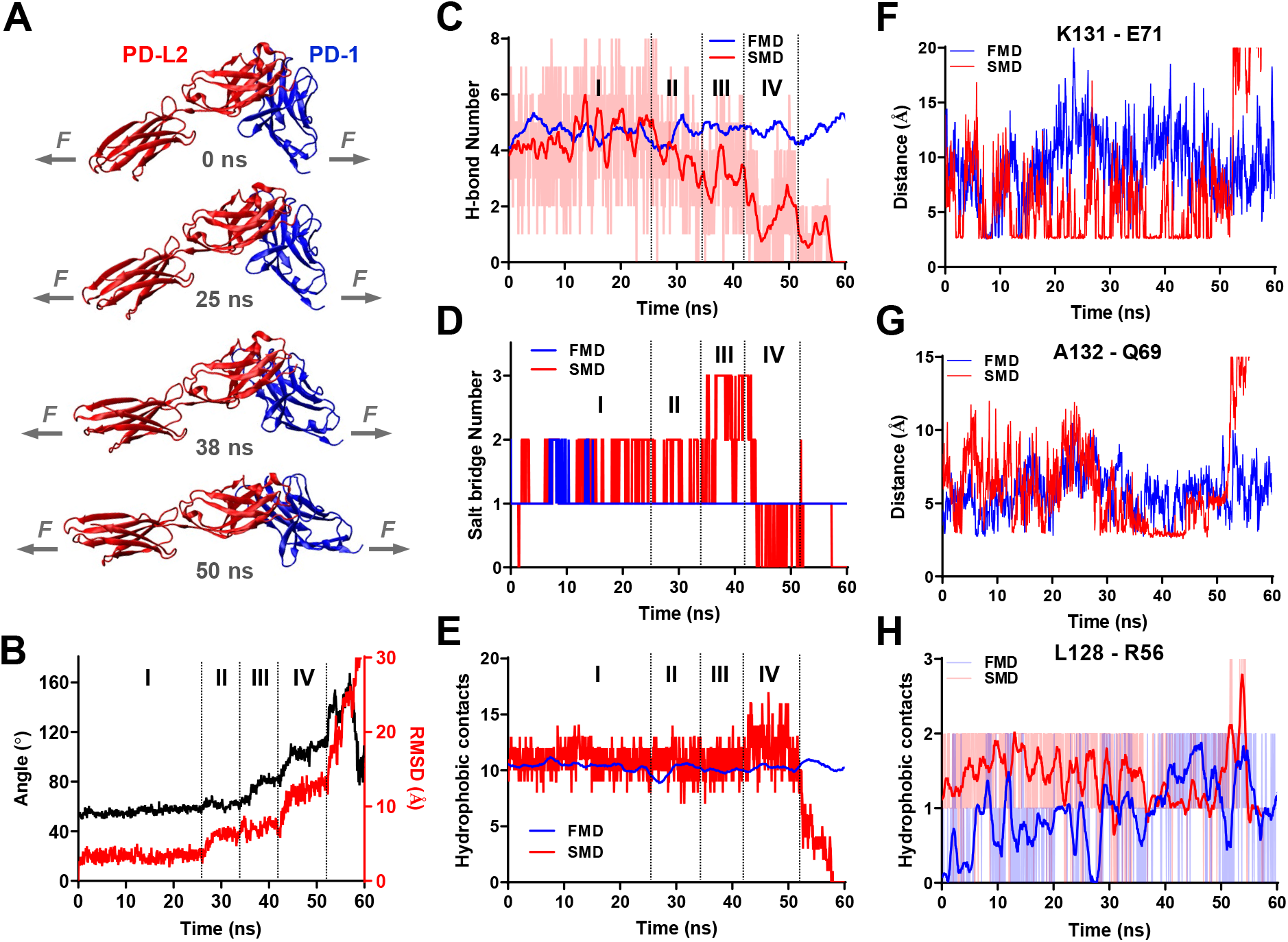
Molecular dynamics (MD) reveals force induced PD-1–PD-L2 conformational change and formation of new atomic-level contacts. (A) Snapshots of PD-1–PD-L2 complex undergoing conformational changes in response to force. (B) Changes in relative angle (black curve, left *y*-axis) and root mean square displacement (RMSD, red curve, right *y*-axis) between PD-1 and PD-L2 in response to force. (C-E) Comparison of total number of hydrogen bond (H-bond, C), salt bridge (D), and hydrophobic contacts (E) between PD-1 and PD-L2 during free MD (FMD) and force steered MD (SMD). (F-H) Comparison of dynamics of possible interactions between indicated residues of PD-1 and PD-L2 during FMD (blue) and SMD (red). Atomic-level contacts were defined by an interatomic distance of < 3.5 Å, which were more frequently observed in FMD than in FMD.

Detailed analysis of bonding events at atomic scale reveals significant force-induced changes of different types of interactions underlying the binding interface rearrangement in SMD, which was absent in FMD. During SMD the total number of hydrogen bond (H-bond) was stable in phase I and shifted to an overall down trend in phase II with brief but significant pull backs in phases III and IV (Fig. 4C). Salt bridge interactions were enhanced by force in late phase I through phase III before being suppressed in phase IV (Fig. 4D). In contrast, total number of hydrophobic contacts remains unchanged until increases by ∼50% in phase IV (Fig. 4E). Together, these observations indicate that unlike spontaneous dissociation in the absence of force where all atomic interactions holding the PD-1–PD-L2 molecular complex together were simultaneously broken, applied force induced formation of different types of new noncovalent contacts while disrupting the original ones, seemingly rendering reinforcement during certain phases of the dissociation when combining all atomic interactions together. In particular, we noticed that some of the force enhanced atomic contacts were not located in the binding pocket or disrupt force-free PD-1–PD-L2 binding when mutated^15^, such as Leu128, Lys131 and Ala132 located in the FG loop of PD-1 (Figs. 4F-H), allowing us to hypothesize that one of their roles is to stabilize PD-1–PD-L2 bond under force.

### PD-1 mutants targeting force-induced atomic contacts impairs PD-1–PD-L2 mechanical stability

To test the aforementioned hypothesis, we made single- and double-residue PD-1 mutations K131A, L128A/K131A, and A132K (Figs. 4F-H) and expressed these mutants on CHO cells at similar levels (Fig. 5A) to test whether eliminating the atomic interactions formed by these residues with PD-L2 under force would abolish the force-enhanced PD-1–PD-L2 bond stability. When analyzed against RBCs coated with PD-L2, all three PD-1 mutants show similar two-dimensional (2D) effective affinity as WT PD-1 (Fig. 5B), consistent with previous studies showing similar PD-L2 Ig staining^15^. However, dynamic force spectroscopic analysis with BFP showed that all three mutants left-shifted the cumulative frequency curves towards lower forces relative to the WT curve (Fig. 5C). Also, the two single-mutants down-shifted the bond lifetime *vs* force curves towards shorter lifetimes relative to the WT curve, and the double-mutant completely converted the catch bond to a slip bond (Fig. 5D). Evaluating the combined effect of ruptured bonds (rupture events) and unruptured bonds (lifetime events) as multiplying bond lifetime by the probability of bond survival at the corresponding force level (1 – cumulative frequency of bond rupture), we estimated the average effective bond lifetime as what would be sampled by PD-1 on cells at given force levels, which modified the bond lifetime *vs* force curves quantitatively but not qualitatively (Fig. 5E). Together, these results have supported our hypothesis and revealed the structural mechanisms underlying the mechanical stability of the PD-1–PD-L2 bond under force.

**Fig. 5.**
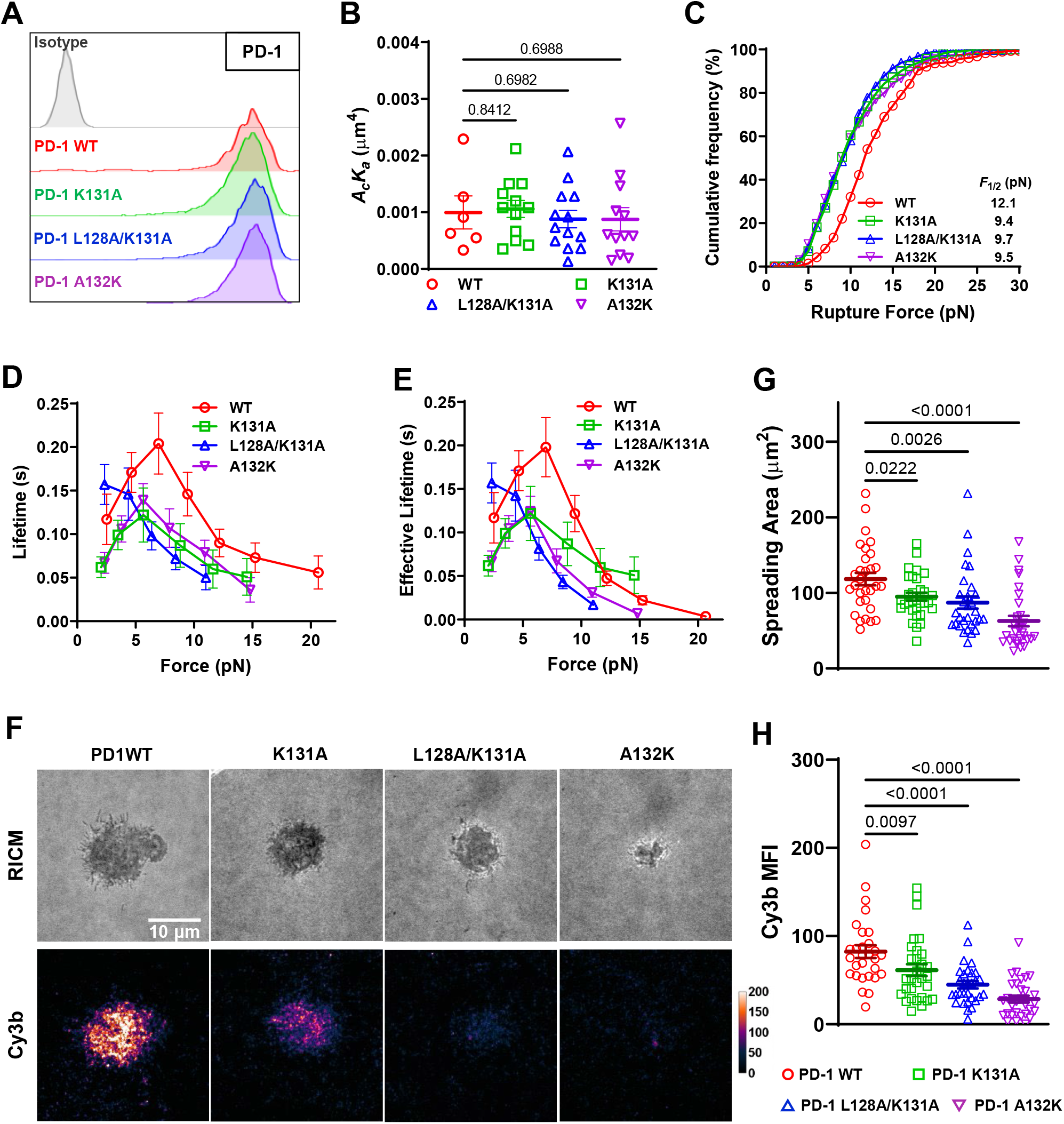
PD-1 mutants preventing force-induced atomic contacts impairs PD-1–PD-L2 mechanical stability. (A) Flow cytometry histograms comparing PD-1 staining of CHO cells expressing WT or indicated mutants of PD-1. (B) 2D effective affinity of PD-L2 binding to CHO cells expressing WT or indicated mutants of PD-1. n = 6, 12, 13, and 12 cell pairs for WT, K131A, L128A/K131A, and A132K, respectively. (C) Cumulative frequencies of rupture force events for PD-L2 binding to PD-1 WT (n = 278 events), K131A (n = 210 events), L128A/K131A (n = 345 events), and A132K (n = 270 events) bonds. p < 0.0001 comparing *F*_1/2_ of WT and each mutant using two-tailed Mann-Whitney test. (D) Mean ± sem bond lifetime vs force plots for single PD-L2 bonds with PD-1 WT (n = 785 events), K131A (n = 625 events), L128A/K131A (n = 759 events), and A132K (n = 780 events). p < 0.0001 comparing lifetime vs force distributions of WT and each mutant using two-tailed two-dimensional Kolmogorov-Smirnov test. (F) Representative RICM and Cy3b fluorescence images of CHO cells expressing PD-1 WT or indicated mutants 30 min after landing on glass surface functionalized with PD-L2-coupled MTP of 4.7 pN threshold force. (G-H) Quantification of cell spreading area (G) and tension signal (H) for conditions in F. n = 29, 30, 30, and 30 pooled from 3 independent experiments. Numbers on graphs represent p values calculated from two-tailed student t-test.

To directly visualize the effect of these mutations on PD-1–PD-L2 bonds under active cellular forces, we analyzed the cell spreading and tension using PD-L2 coupled MTP of 4.7 pN threshold force. Consistent with impaired bond strength and lifetime, both cell spreading and tension signal was reduced in CHO cells expressing these mutants with the level of impairment ranking as K131A < L128A/K131A < A132K (Fig. 5F-H). Since the 2D affinity of the mutants are similar to that of WT, such reductions in spreading and tension indicate that the reduced stability of bonds between PD-L2 and PD-1 mutants has decreased the numbers of opened MTPs or become less able to support cell spreading.

### PD-1 mutants with impaired PD-1–PD-L2 mechanical stability show reduced inhibitory function

Combining the data of Figs. 1&2 prompt us to hypothesize that the inhibitory function of PD-1 requires mechanical support of PD-Ligand to counter-balance the force the cell exerts on the PD-1 bonds. To test this hypothesis, we investigated whether disrupting PD-1–PD-L2 mechanical stability would affect PD-1 function. We expressed PD-1 K131A, L128A/K131A, and A132K mutants in NFκB::eGFP reporter Jurkat cells to similar level as WT PD-1 (Fig. 6A) and tested their ability to suppress TCR-CD3 triggered GFP induction by coculturing them with TSC cells with or without PD-L2. Comparing to control TSC not expressing PD-L2 (TSC-CTRL), TSC-PD-L2 cells triggered robust suppression of GFP induction in reporter cells expressing WT PD-1 relative to plain cells not expressing PD-1 (Fig. 6B-D). The inhibitory effect was significantly reduced by all three PD-1 mutants with the level of impairment ranking as K131A < L128A/K131A < A132K (Figs. 6B-D), which are the same in cell spreading and tension measurements (Figs. 5F-H). Together, these data suggest that mechanical forces on PD-1–PD-Ligand bonds critically regulate PD-1 ligand interaction and function.

**Fig. 6.**
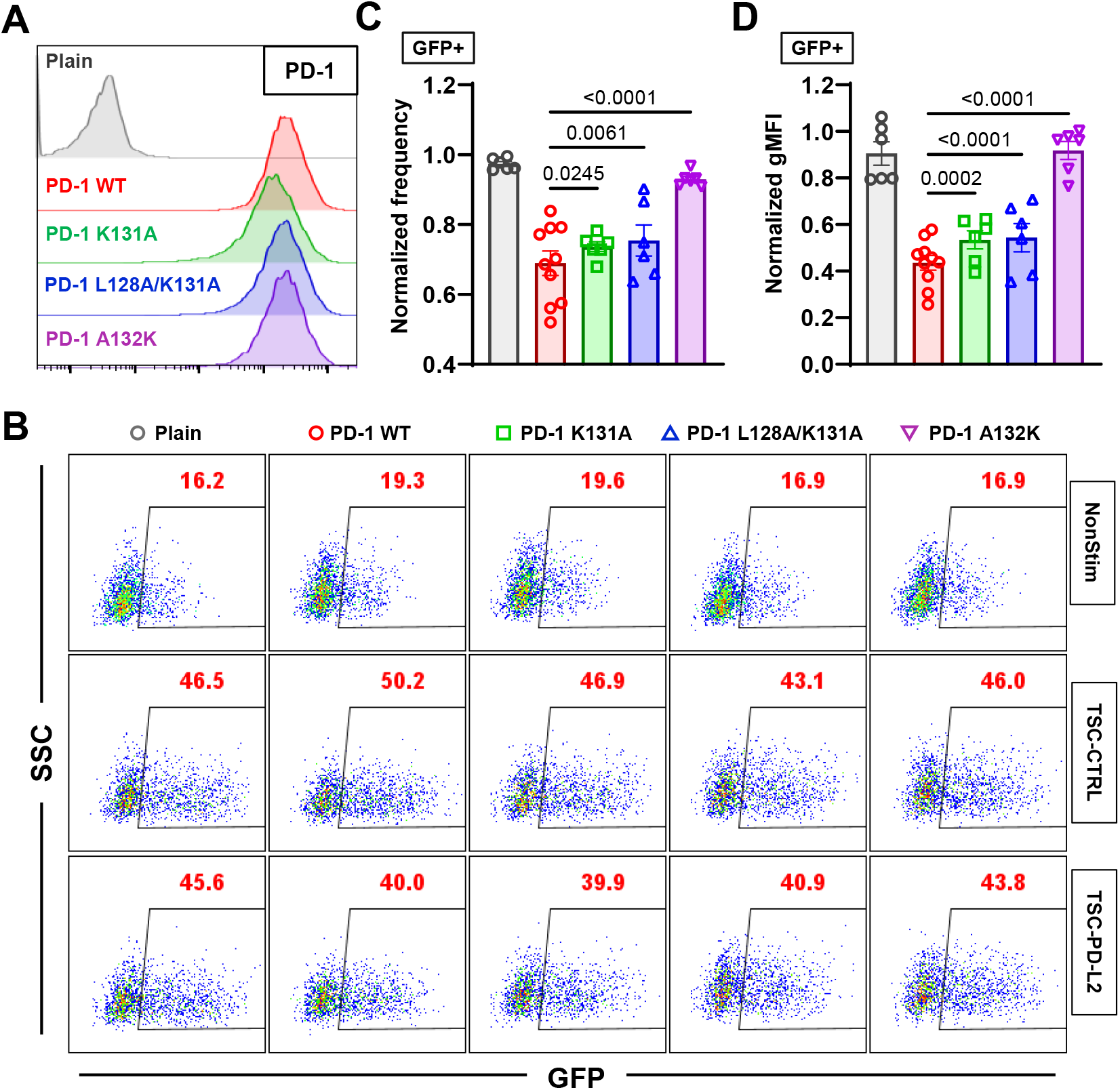
PD-1 mutants with impaired PD-1–PD-L2 mechanical stability demonstrate reduced inhibitory function. (A) Flow cytometry histograms comparing PD-1 staining of NFκB::eGFP reporter Jurkat cells expressing no, WT, or indicated mutants of PD-1. (B) Representative SSC vs GFP plots of reporter Jurkat cells 24 hr after stimulation with indicated conditions. (C-D) Quantification of GFP expression for conditions in B. Normalized frequency (C&G) and normalized geometric mean fluorescence intensity (gMFI) (D&H) were calculated as (sample – averaged background)/(anti-CD3 control – averaged background). n = 6 for plain, PD-1 K131A, L128K/K131A, and A132K pooled from 3 independent experiments or n = 10 for PD-1 reporter cells pooled from 5 independent experiments. Numbers on graphs represent p values calculated from two-tailed student t-test.

## Discussion

Despite the success of immune checkpoint blockade of PD-1 or PD-L1 in cancer immunotherapy, the molecular mechanism of PD-1 triggering as to how ligand binding translates into biochemical signaling remains poorly understood. Dissecting the structural, biophysical, and biochemical bases of this mechanism would also expand our fundamental understanding of signal initiation of a general class of transmembrane receptors and guide the design of therapeutic agents targeting these receptors. While formation of PD-1 microclusters after ligand binding and their co-localization with TCR and CD28 were observed and found critical for PD-1’s inhibitory function, other studies suggest that such co-localization is not required as HEK cells expressing PD-L1 is able to suppress the activating signal of bead-coated anti-CD3 and anti-CD28 on T cells in a co-culture system^11^. Consistent with the latter finding, our observation that bead-coated PD-Ligands were able to suppress soluble anti-CD3 induced NFκB activation also support the contention that localized PD-1 signaling is able to suppress TCR-CD3 signaling globally. A deep dive into the triggering process also begs the question as to whether the microcluster formation and co-localization drive PD-1 signal initiation or instead are merely their consequence. What seemingly have made this more perplexing are the findings that soluble isoforms of PD-L1 or PD-L2 in patients of different cancers fail to suppress T cell function *in vitro* and *in vivo* despite their ability to bind PD-1^9-11,30,31^. Comparing the same ectodomain of PD-L1 and PD-L2 in the form of soluble tetramer to immobilized on bead surface, we further confirmed that surface anchor of the PD-Ligand is required to induce PD-1 function, an important requirement that has been largely ignored. Moreover, the failure of multimeric ligand binding to trigger PD-1 signaling suggests that crosslinking of PD-1 by soluble PD-Ligand is not enough to initiate its signaling, and the formation of microclusters are likely a consequence of ligand binding (and signaling) instead of a mechanism of triggering.

What, then, does the bead-coated PD-Ligand provide but soluble PD-Ligand cannot? A recent study revealed activated T cells actively apply mechanical forces on PD-1, which were detected using DNA-based MTP presenting either PD-L2 or anti-PD-1^13^. Interestingly, such forces could originate from cell-intrinsic mechanical sampling processes as we also observed cellular forces on PD-1–PD-L1 or PD-1–PD-L2 bonds when expressing PD-1 in CHO cells. In support of this hypothesis, PD-1 expressing CHO cells are able to form “synapse” like interface with PD-L2 expressing CHO cells with accumulation of the receptors and ligands within the interface^15^. This mechanical sampling process is quite intriguing because it suggests that the cell is actively probing the ON or OFF state of its membrane PD-1 molecules by exerting force on them, and only those bound to an anchored ligand with mechanical support to counter-balance such forces respond to this probing and are recognized as in the ON state, whereas those without ligand binding or bound to soluble ligand cannot be properly detected and thus remain in the OFF state. In other words, successful PD-1 triggering requires not only the formation of PD-1–PD-Ligand bonds, but also for these bonds to respond mechanically so as to be detected by the cells as a productive binding event. Structurally, this is also plausible, because there could hardly be any ON/OFF conformations defined from a molecule as simple as PD-1: a single IgV domain linked to intracellular domain by ∼20 AA peptide of stalk and transmembrane region. In support of this hypothesis, we found that dampening the mechanical support of PD-Ligands by extending their length reduced the capacity of PD-1 to trigger function without affecting its PD-1 binding.

It follows from the above mechanical sampling hypothesis that the efficiency of PD-1 triggering is not merely determined by the PD-1 affinity for ligand as measured under force-free condition. In addition, the mechanical stability of the PD-1–PD-Ligand bonds – how the complex responds to force – would also be critical to the triggering process. This suggests bonds that readily rupture under force are less effective in initiating signals, whereas bonds that are more mechanically stable under force are more effective in triggering PD-1, implying the ability of PD-1 to sense the stiffness of the PD-Ligand expressing cell. Interestingly, we observed catch bonds for both PD-1–PD-L1 and PD-1–PD-L2, a phenomenon where increasing forces below an optimal level (7 – 8 pN for PD-1) promote bond stability instead of accelerating dissociation. This force-induced reinforcement of bond sustainability, or mechanical stability, is more profound for PD-1–PD-L2 than PD-1–PD-L1 according to the rupture force distribution, catch bond profile, and MTP tension signal, suggesting a potential mechanism for ligand discrimination by PD-1. These results, although phenotypically similar to that observed in TCR antigen recognition, differ in its underlying structural mechanisms. Our SMD simulations of PD-1–PD-L2 dissociation reveal that the unique positioning of the two IgV domains from PD-1 and PD-Ligand with “side-to-side” binding using relatively flat β-sheets makes the complex very sensitive to mechanical loads. Forces applied to the two C-termini of the complex readily generate a “peeling” effect. In response to force, the two IgV domains rotate and translate relative to each other, transitioning into a stretched and aligned conformation. More importantly, such large-scale rearrangement of the complex is coupled with formation of new atomic-level interactions, which were not observed under force-free FMD conditions. These *in silico* studies and the experimentally measured catch bond profiles suggest that mechanical stability of PD-1–PD-Ligand is required by the mechanical sampling process of PD-1, without which PD-1 triggering may also fail as seen in the case of soluble ligand completely loosing mechanical support.

Indeed, comparing SMD *vs* FMD results we identified residues of PD-1 that were not in contact with PD-L2 in the crystal structure and in FMD but formed new interactions in SMD. Mutations aiming to prevent the formation of these new interactions did not alter the force-free PD-1–PD-L2 2D affinity, consistent with the assertion that they do not contribute to static binding, but significantly reduced the bond stability under force, as shown by lower rupture force, shorter and altered profile of bond lifetime, as well as reduced MTP tension signal. Consequently, these mutants demonstrate impaired ability to trigger PD-1 function with the same rank-order as that in the tension signal. These results indicate force and mechanical stability of PD-1–PD-Ligand bonds play critical role in the triggering of PD-1 signaling and function.

Overall, our data suggest a potential PD-1 triggering mechanism consisting of three key components: 1) mechanical sampling: cells actively apply forces on PD-1 to probe ligand binding; 2) mechanical support: pulling force from the PD-Ligand anchoring surface counter-balances the pulling force from PD-1; 3) mechanical stability: forces on PD-1–PD-Ligand bonds modulate their dissociation kinetics and triggering efficiency. Formation and movement of PD-1 microclusters upon ligand binding may be subsequent steps following such recognition event and provide mechanical feedback as PD-1 is being actively transported. Mechanistically, it is of great interest as to how mechanical information is integrated into this recognition process and finally leads to phosphorylation of PD-1. On the translational side, our results suggest modulating the mechanical support and mechanical stability of the PD-1–PD-Ligand system as a potential strategy to regulate PD-1 agonism for disease treatment.

## Methods

### Proteins and antibodies

Mouse PD-L1 and PD-L2 with C-terminal biotin produced in CHO cells^43^ were generous gifts of Dr. Simon J. Davis (University of Oxford, United Kingdom). PE-anti-PD-1 (clone RMP1-30, 1:20) and its isotype control PE-Rat IgG2b,κ (clone RTK4530, 1:20) were purchased from Biolegend. PE-anti-PD-L1 (clone MIH5, 1:20) and its isotype control PE-Rat IgG2a,λ (clone B39-4, 1:20) together with PE-anti-PD-L2 (clone TY25, 1:20) and its isotype control PE-Rat IgG2a,κ (clone R35-95, 1:20) were purchased from BD Biosciences. APC-anti-CD45.2 (clone 104, 1:100) was purchased from Biolegend. Purified anti-CD3 (clone OKT3) and anti-mouse IgG2a (clone RMG2a-62) were purchased from Biolegend. Purified antiPD-1 blocking antibody (clone 29F.1A12), biotinylated antiPD-1 antibodies (clone 29F.1A12 and clone RMP1-30), and biotinylated isotype controls (Rat IgG2a,κ clone RTK2758 and Rat IgG2b,κ clone RTK4530) were purchased from Biolegend. Biotinylated antiPD-1 antibody (clone J43) and isotype control (clone eBio199Arm) were purchased from Invitrogen.

### Cells

NFκB::eGFP reporter Jurkat cells and T-cell stimulator cells (TSC) expressing a membrane-anchored scFv of antiCD3 (clone OKT3) were generous gifts of Dr. Peter Steinberg (Medical University of Vienna, Austria). Jurkat cells, TSC, and CHO cells (ATCC) were cultured in RPMI 1640 supplemented with 10% FBS, 100 U/mL penicillin, 100 μg/mL streptomycin, 2 mM L-glutamine, and 20 mM HEPES. HEK 293T cells (ATCC) were cultured in DMEM supplemented with 10% FBS, 6mM L-glutamine, 0.1 mM MEM non-essential amino acids, and 1 mM sodium pyruvate. Human RBCs were isolated from healthy donors and used following previously described protocols ^43^ approved by the Institutional Review Board of the Georgia Institute of Technology.

### Overexpression of PD-1 and PD-Ligands in CHO cells, Jurkat cells and TSC

All PD-1 mutants were generated using Q5 site-directed mutagenesis kit (NEB) following the manufacture’s protocol. To generate CHO cells expressing WT or MT mouse PD-1, cells were transfected with pcDNA3.1 encoding full-length mouse PD-1 or its mutant using nucleofection (Lonza). Transfected cells were sorted for uniform PD-1 staining and culture in medium with 0.4 cells and reporter Jurkat cells stably expressing WT or MT chimeric PD-1 (mhPD-1) consisting of mouse PD-1 ectodomain (Met1 to Met169) and human PD-1 transmembrane and intracellular domain (Val171 to Leu288), full-length mhPD-1 were subcloned in lentiviral vector pLenti6.3. Lentivirus were produced by transfecting HEK 293T cells with a mixture of mhPD-1-pLenti6.3, pMD2.G (Addgene #12259), and psPAX2 (Addgene #12260) using Lipofectamine 3000 (Thermo Fisher Scientific) following manufacture’s protocol. CHO cells and reporter Jurkat cells were transduced overnight with 1:1 mixture of culture medium and lentiviral supernatant. Cells were then sorted for uniform and similar PD-1 expression across WT and MT. To generate TSC expressing mouse PD-L1 or PD-L2, full-length PD-L1 or PD-L2 were subcloned into pMSCV-IRES-Thy1.1 (pMIT1.1) vector. Retrovirus were produced by transfecting HEK 293T cells with a mixture of PD-L1-pMIT1.1 (or PD-L2-pMIT1.1) and pCL-Eco (Addgene #12371) using Lipofectamine 3000 (Thermo Fisher Scientific) following manufacture’s protocol. TSC were transduced by spinoculation on retronectin-coated plate (Takara) with 1:1 mixture of culture medium and retroviral supernatant. Cells were subjected to repeated rounds of transduction and sorting to get desired PD-1 ligand expression.

### Flow cytometry

Cells were stained in 100 μl of FACS buffer (PBS supplemented with 5mM EDTA and 2% FBS) containing fluorescently labeled antibodies (dilutions indicated above) for 30 min at 4 °C. After staining cells were washed twice with FACS buffer and analyzed using Fortessa flow cytometer (BD Biosciences). Flow cytometric data were analyzed using Flowjo (TreeStar). Cells were gated based on FSC vs SSC for mono-population where expression of PD-1 and PD-Ligands were stained. In Jurkat and TSC coculture experiments, Jurkat cells were gated on CD45.2-population to exclude TSC.

### Stimulation of Jurkat cells

For experiments illustrated in Fig. 1A, 50,000 NFκB::eGFP reporter Jurkat cells were co-cultured with 50,000 TSC-CTRL, TSC-PD-L1, or TSC-PD-L2 for 24 hours. After stimulation, cell mixture was stained with APC-antiCD45.2, washed, and analyzed by flow cytometry. For experiments illustrated in Figs. 1E, 1F and S2, 50,000 NFκB::eGFP reporter Jurkat cells were stimulated with 10 μg/ml anti-CD3 (clone OKT3) and 5 μg/ml anti-mouse IgG2a,κ secondary antibody for 24-30 hrs. For soluble PD-Ligand (or antibody) groups, a final concentration of 20 μg/ml pre-made PD-Ligand (or antiPD-1 antibody or isotype Ig) tetramer (concentration excluding SA) or the corresponding amount of SA were added. For bead-coated PD-Ligand (or antibody) groups, 3 μl of SA beads (Dynabeads M-280 Streptavidin) were coated with 300 ng of PD-Ligand (or antibody or isotype Ig) for 1 hr at room temperature. After coating, beads were washed twice with PBS and mixed with cells at 10:1 bead-to-cell ratio. After stimulation, cells were washed and resuspend in FACS buffer for flow cytometric analysis. For experiments illustrated in Fig. 1I, cells were stimulated with the same conditions as in Fig. 1F with 10:1 bead-to-cell ratio except for using glass beads that were activated with MPTMS (Sigma) followed by conjugation with SA using SMCC or SM[PEG]_24_ crosslinker (Sigma). Normalized frequency and normalized geometric mean fluorescence intensity (gMFI) of GFP+ population were calculated as (sample – averaged background)/(anti-CD3 control – averaged background).

### DNA hairpin sequences

**A21B-Cy3B:** Cy3B – CGC ATC TGT GCG GTA TTT CAC TTT - /3Bio/ **SH-BHQ2:** /5-ThioC6-5/ - TTT GCT GGG CTA CGT GGC GCT CTT - /3BHQ_2/ **12 pN HP:** GTG AAA TAC CGC ACA GAT GCG TTT GGG TTA ACA TCT AGA TTC TATTTT TAG AAT CTA GAT GTT AAC CCT TTA AGA GCG CCA CGT AGC CCA GC **4.7 pN HP:** GTG AAA TAC CGC ACA GAT GCG TTT GTA TAA ATG TTT TTT TCA TTT ATA CTTTAA GAG CGC CAC GTA GCC CAG C

### AuNP DNA tension sensor preparation

AuNP-based tension probes were prepared following our previous work^13,34,44^. No. 2 glass coverslips (VWR: 48382085) were sonicated for ∼5 minutes in nanopure water (18.2 MΩ) followed by ∼5 minutes of sonication in pure ethanol. Coverslips were dried at 80°C for 10 minutes and then etched in a fresh piranha solution containing 37.5% v/v hydrogen peroxide (30% solution) and 62.5% v/v concentrated sulfuric acid. Coverslips were then washed six successive times in nanopure water, followed by three successive washes in pure ethanol. Coverslips were then immersed in a 3%v/v APTES (Sigma-Aldrich: 440140) solution in ethanol for one hour at room temperature. Following this reaction, coverslips were washed three times with ethanol and baked in an 80°C oven for 30 minutes. After cooling, the silanized coverslips were incubated with 1% w/v lipoic acid polyethylene glycol-succinimidyl ester (Biochempeg: HE039023-3.4K, MW 3400) and 10% w/v monofunctional polyethylene glycol-succinimidyl ester (Biochempeg: MF001023-2K, MW 2000) in a freshly prepared 0.1M NaHCO33 solution for at least 1 reacted hour at room temperature. Following PEG functionalization, coverslips were washed with nanopure water and blocked with a 1% w/v solution of Sulfo-NHS-Acetate (Thermo: 26777) in a freshly prepared 0.1M NaHCO33 solution for 30 minutes to neutralize the positive charges of the amines and prevent nonspecific DNA binding to the surface. Following blocking, coverslips were washed with nanopure water and incubated with 400 μL of a 20nM solution of 8.8 nm gold nanoparticles (AuNPs, nanoComposix) for 30 minutes. Finally, the DNA tension probe hairpins were assembled in 1M NaCl by mixing Cy3B-labeled A21B (0.33 μM), BHQ2 strand (0.33 μM), and hairpin strand (0.3 μM) in a 1.1: 1.1: 1 ratio. The solution was heated to 95°C and cooled to 25°C in a thermocycler over a period of 30 minutes. After heating and cooling, 2.7 μM of passivating BHQ2 ssDNA was added to the DNA hairpin solution. Following DNA assembly, AuNPs were rinsed off the coverslips with nanopure water followed by rinsing with a 1M NaCl solution. Coverslips were placed in a petri dish and 100 μL of the DNA hairpin was added to the surface. A second AuNP-functionalized surface was placed upside-down on top of the first coverslip to create a “sandwich.” Sandwiched slides were incubated with DNA overnight at 4°C and further functionalized the following day.

### DNA hairpin ligand functionalization

DNA sandwiched coverslips were removed from 4°C and separated into single coverslips. Coverslips were washed in 1X PBS and 200 uL of 40 μg/mL streptavidin (Rockland: S000-01) solution was added to each. Streptavidin was incubated on the coverslips for 1 hour. Following incubation, coverslips were washed with 1X PBS and further functionalized with C-terminal-biotinylated mPDL1 or mPDL2 at a concentration of 15 μg/mL and placed in imaging chambers containing 1 mL of cellular imaging media (1x HBSS with 10 mM HEPES).

### Fluorescence Imaging

Imaging was done on a Nikon Eclipse Ti microscope, operated by Nikon Elements software, a 1.49 NA CFI Apo 100x objective, perfect focus system, and a TIRF laser launch with a 80mW 561nm laser. A reflection interference contrast microscopy (RICM) cube (Nikon: 97270) was used for imaging. An X-Cite 120 lamp (Excelitas) was used for widefield epifluorescence illumination. Images were acquired with an Andor iXon Ultra 897 electron-multiplying charge-coupled device. Fluorescent images acquired with TIRF excitation were taken with 100 ms exposure time, 10% laser power, and no gain.

### Micropipette adhesion frequency assay

As previously described^37,43,45,46^, the 2D effective affinity between PD-L2 and WT or MT PD-1 were measured by analyzing the bond formation between PD-L2-coated human RBCs and target cells expressing WT or MT PD-1. In brief, RBCs were biotinylated, coated with SA, washed, and then coated with biotinylated PD-L2. During experiments, a PD-1 expressing cell was repeatedly brought into contact with a PD-L2-coated RBC, held for 5 seconds, and then separated. Due to the ultrasoft spring constant of the RBC membrane, PD-1–PD-L2 bonds formed during the contact (and last till separation) caused stretch of RBC membrane upon separation, and therefore defined as an “adhesion” event. The process was repeated for 30 – 50 cycles per RBC-Target cell pair and an averaged adhesion frequency (*P*_a_) was calculated. *P*_a_ is related to the 2D effective affinity of the molecular interaction in question according to the following equations:

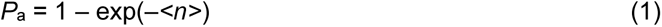

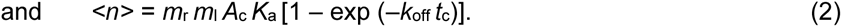

Here *<N>* is the average number of bonds per contact, *m*_r_ and *m*_l_ are the respective densities of PD-1 on target cell and PD-L2 on RBC, *A*_c_ is contact area (in μm^2^), *K*_a_ is 2D affinity (in μm^2^), and *k*_off_ is off-rate (in s^-1^). With long contact duration such as 5 s used in these experiments, *k*_off_ *t*_c_ >> 1^37^, *P*_a_ and <*n*> approach equilibrium, and the 2D effective affinity *A*_c_*K*_a_ was estimated by normalizing <*n*> against *m*_r_ and *m*_l_ that were measured using PE-labeled monoclonal antibody together with QuantiBRITE PE standard beads (BD Biosciences):

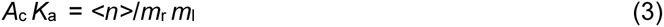

### Biomembrane force probe

A previously described^20,43^ BFP was used to measure the rupture force and bond lifetime at certain clamp force levels of PD-1–PD-Ligand bonds. A target cell expressing PD-1 is repetitively brought into contact with a PD-L1- or PD-L2-coated glass bead attached to a micropipette aspirated RBC. The displacement of the bead is tracked at 1000 fps with nanometer precision and is then translated into force trace with a BFP spring constant preset to 0.3 pN/nm. Compression of RBC during approaching and contact phase generate negative force values, whereas tension on the molecular bond formed between the ligand on the bead and the receptor on the target cell pulls the bead away from its resting position during target cell retraction. For rupture force measurement, the cell was retracted continuously after each contact. If a bond formed the force would increase linearly until bond rupture at which point the force level was recorded as the rupture force value. Hundreds of bond rupture events from repeated cycles were pooled to construct the rupture force histogram and the cumulative frequency of rupture events. For bond lifetime measurement, the target cell was initially retracted and then held at a distance corresponding to the preset clamp force level. The force sustained until the bond ruptured, with the total duration defining the bond lifetime under the corresponding clamped force level. Hundreds of bond lifetime events were pooled from repeated measurements in a range of clamp forces, which were used to construct the bond survival frequencies at different force binds and to calculate average bond lifetime vs average force by binning the events across multiple force levels.

### Molecular dynamics simulation

The crystal structure of mPD-1/mPD-L2 (PDB code: 3BP5) was used as the initial coordinates of the atoms. The protein structure was placed and solvated in a rectangular box of TIP3P water molecules with a 15 Å minimal distance between boundaries and proteins imposed to avoid protein interacting with their mirror images. Water molecules were neutralized by adding Ca^2+^, Cl^-^, and Na^+^ ions to create ∼50 mM calcium concentration and ∼100 mM ionic concentration that are physiologically relevant. The system had in total ∼200,000 atoms (including 1 Ca^2+^, 66 Cl^-^, and 66 Na^+^) in a water box of 160 × 78 × 60 Å^3^. The x-axis was increased to accommodate the steered simulations and warranty that the protein complex was always inside the simulation box to avoid interaction with mirror images. Free molecular dynamics (FMD) and steered molecular dynamics (SMD) simulations^47^ were performed using the NAMD program^48^ and all topology and parameter files were generated using the CHARMM27 all-atom force field for proteins^49^. The SHAKE^50^ was used to constrain bond lengths involving bonds to hydrogen atoms. Periodical boundary condition was used along with particle mesh Ewald method for electrostatic interaction with a grid spacing of 1.0 Å and a 12-Å cutoff for van der Waals interaction. The time-step employed in all simulation was 2 fs with a 12 Å non-bonded cutoff and with long-range non-bonded interactions evaluated every two steps. The systems were initially energy-minimized with a conjugate gradient method for three stages of 50,000 steps each: all atoms of the proteins fixed, then only backbone atoms fixed, and finally all atoms free. After energy minimization, the system was heated up from 0 to 300 K in 300 ps and then equilibrated for 1 ns with pressure and temperature control. The temperature was held at 300 K using Langevin dynamics and the pressure was held at 1 atm by Langevin piston method. FMD was performed on the equilibrated system for 100 ns to simulate an environment without mechanical force input. SMD was then run using results of FMD simulations as initial conditions to mimic the presence of mechanical forces on the PD-1– PD-L2 interactions. For SMD, the C_α_ atom of Met136 of PD-1 domain was pulled through a spring with a spring constant of 70 pN/Å at a constant speed of 1 Å/ns. The C_α_ atom of Leu229 of PD-L2 was constrained and held fixed to its equilibrated position. Simulation data were recorded at 50 ps steps unless stated otherwise. New interaction formation was defined using the interatomic distance following criteria: a hydrogen bond is defined between an atom that has a hydrogen bond (the Donor) and another atom (the Acceptor) if their distance is less than the cut-off distance (3.2 Å) and the angle Donor-H-Acceptor is less than the cut-off angle (20°); a salt bridge is created if the distance between any of the oxygen atoms of acidic residues and the nitrogen atoms of basic residues is within the cut-off distance (3.5 Å) in at least one frame. The principal moments of inertia of the protein were used to measure the angle between interacting domains.

### Statistical analysis

Statistics comparing the mean values of two groups were calculated based on two-tailed student-*t* test as indicated in figure legends. Statistics comparing two lifetime vs force distributions were calculated based on two-tailed two-dimensional Kolmogorov-Smirnov test.

## Acknowledgments

We thank Simon J. Davis for providing the mouse PD-L1 and PD-L2 with C-terminal biotin produced in CHO cells and Peter Steinberg for providing the NFκB::eGFP reporter Jurkat cells and T-cell stimulator cells (TSC). The MD simulations were supported by an NSF award (MCA08X014) in advanced computing infrastructure for U.S. and performed in the Extreme Science and Engineering Discovery Environment (XSEDE). This work was supported by NIH grants R01CA243486 (to C.Z.) and U01CA250040 (to C.Z. and R.A.) and NIH NCI F31 grant F31CA243502 (A.V.K).

---

**Video S1.** Realtime RICM and Cy3B images of a PD-1 CHO cell spreading on PD-L2-coupled tension probe of 4.7 pN threshold force.

**Video S2.** SMD simulated trajectory for pulling the mPD-1–mPD-L2 complex. The C-terminal residue of mPD-1 was harmonically constrained and the C terminus of mPD-L2 was pulled by a dummy spring moving at ∼0.1 nm/ns with a spring constant of ∼70 pN/nm.

**Fig. S1.**
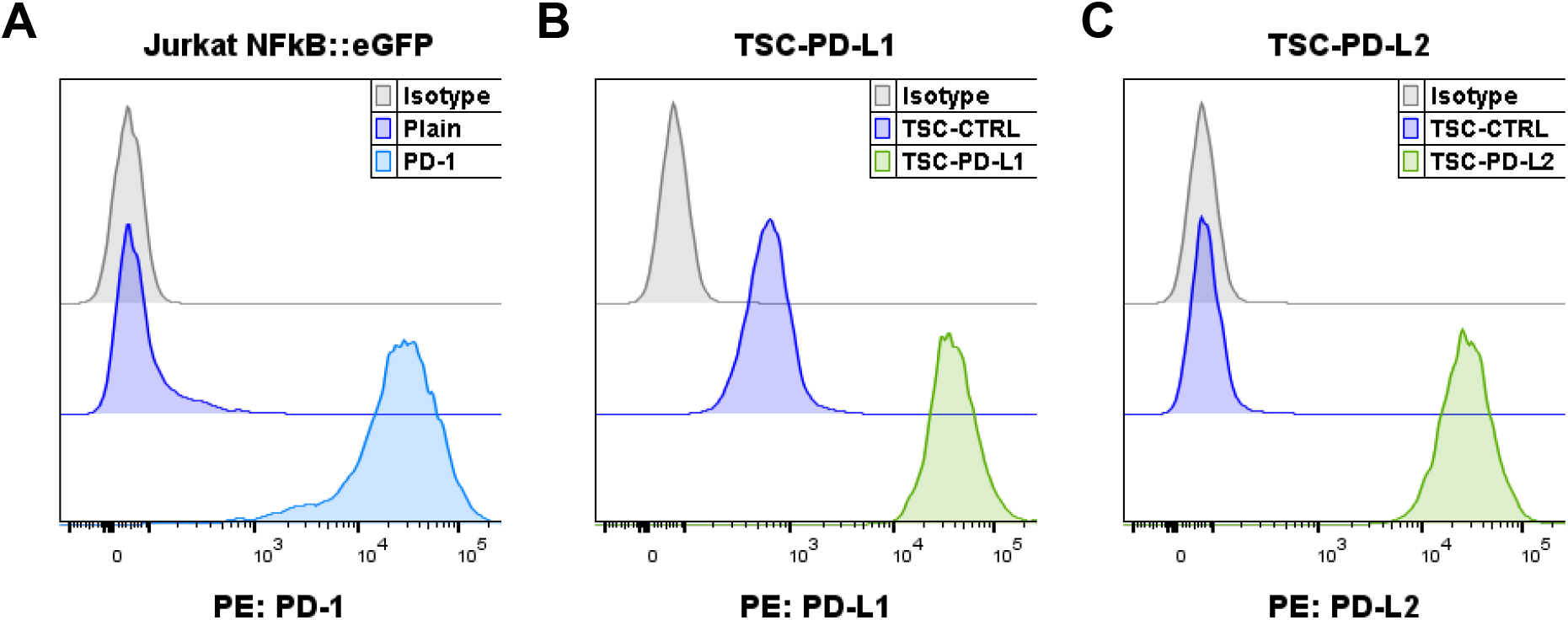
(A) Flow cytometry histograms of PD-1 staining of plain or PD-1 transduced NFκB::eGFP reporter Jurkat cells. (B-C) Flow cytometry histograms of PD-L1 (B) or PD-L2 (C) staining of TSC without transducing PD-1 ligands (TSC-CTRL) or transduced with PD-L1 (TSC-PD-L1) or PD-L2 (TSC-PD-L2).

**Fig. S2.**
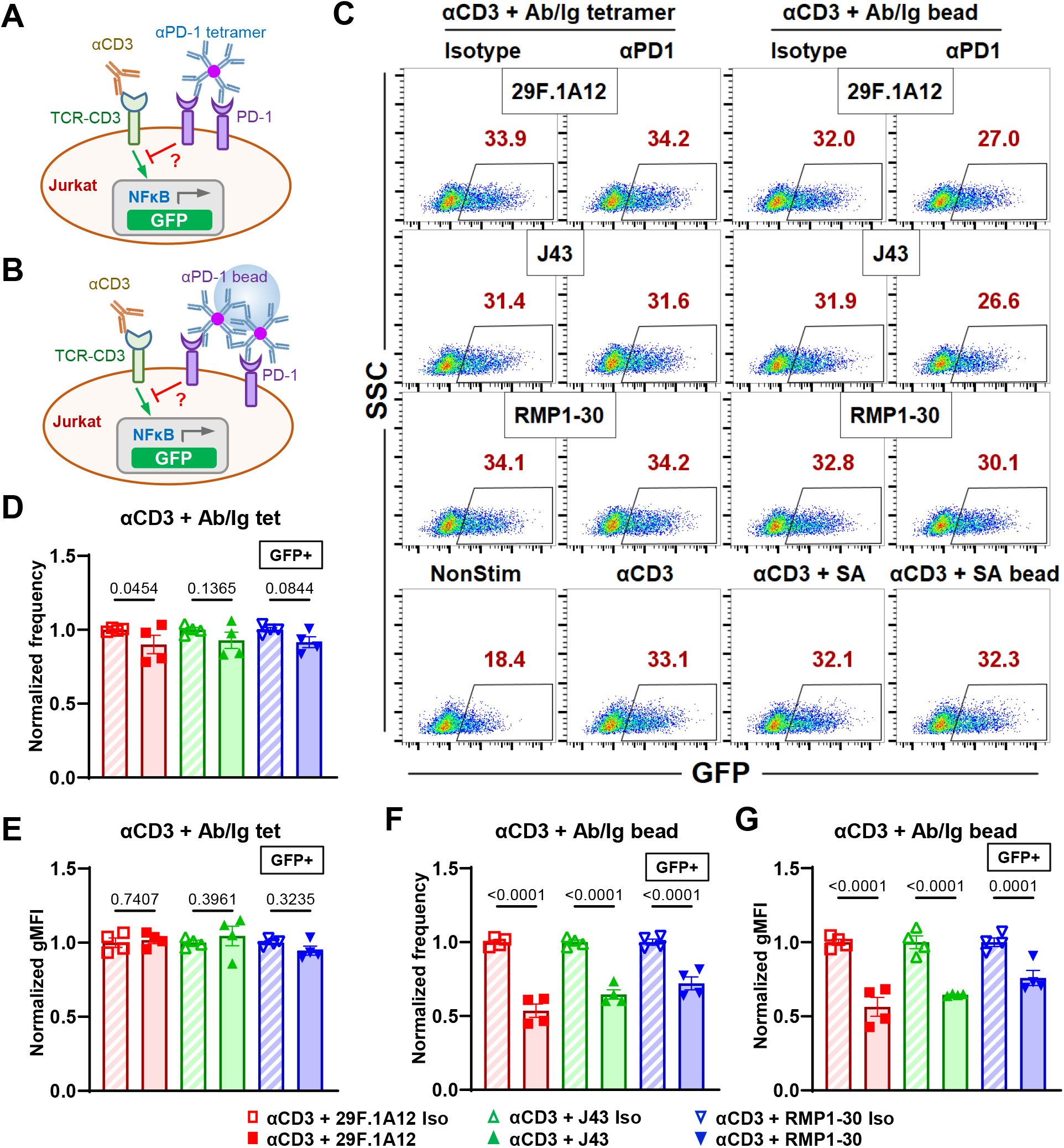
Immobilized anti-PD-1 antibody triggers PD-1 function. (A-B) Schematics of stimulating NFκB::eGFP reporter Jurkat cells with soluble anti-CD3 and soluble (A) or bead-coated (B) anti-PD-1 antibody (or isotype Ig). (C) Representative SSC *vs* GFP plots of reporter Jurkat cells 24 hr after stimulation with indicated conditions. (D-G) Quantification of GFP expression for conditions in A. n = 4 for all conditions pooled from two independent experiments.

**Fig. S3.**
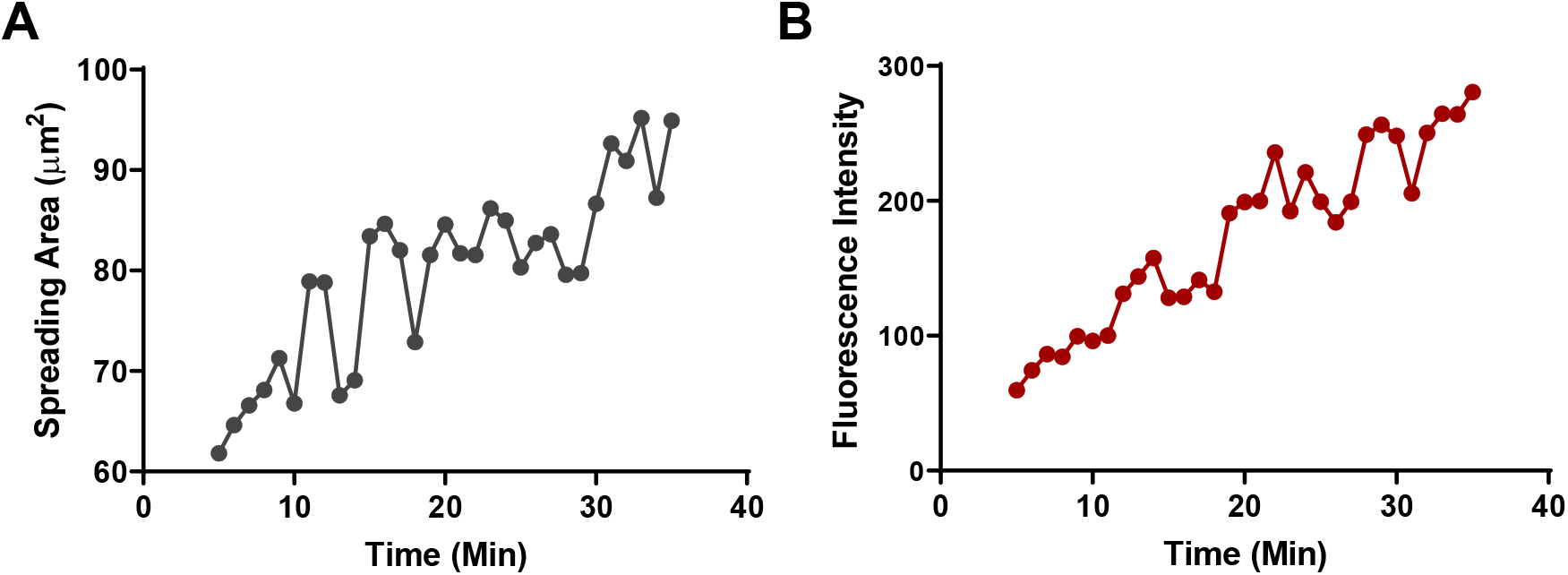
Realtime quantification of spreading area (A) and tension signal (B) of a PD-1 CHO cell spreading on PD-L2-coupled MTP of 4.7 pN threshold force as shown in Video S1.

**Fig. S4.**
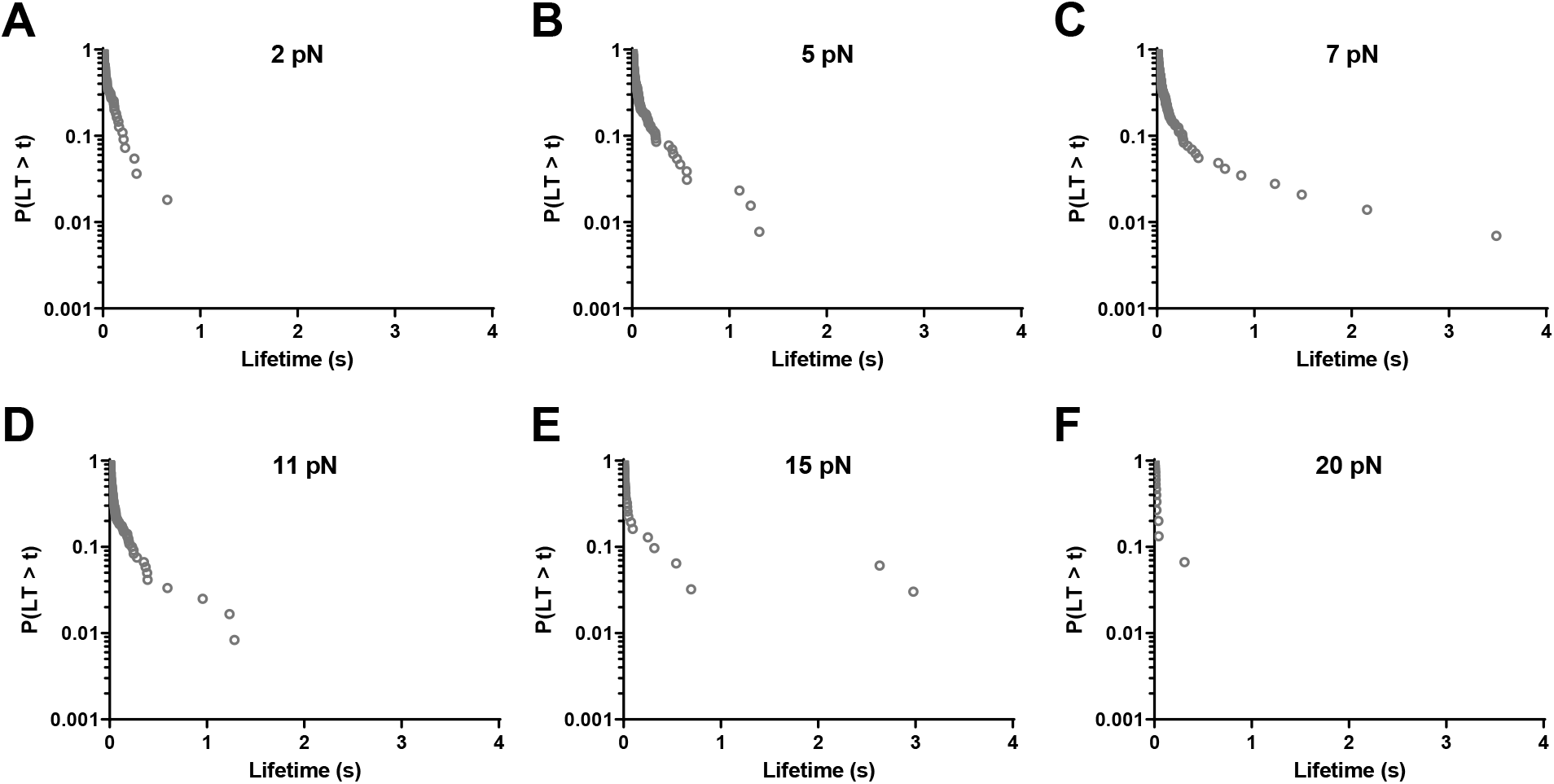
Survival frequencies (fraction of event with lifetime greater than *t*) of PD-1–PD-L1 bond lifetime for different force bins. n = 55, 129, 144, 120, 33, and 16 lifetime events for (A) to (F) respectively.

**Fig. S5.**
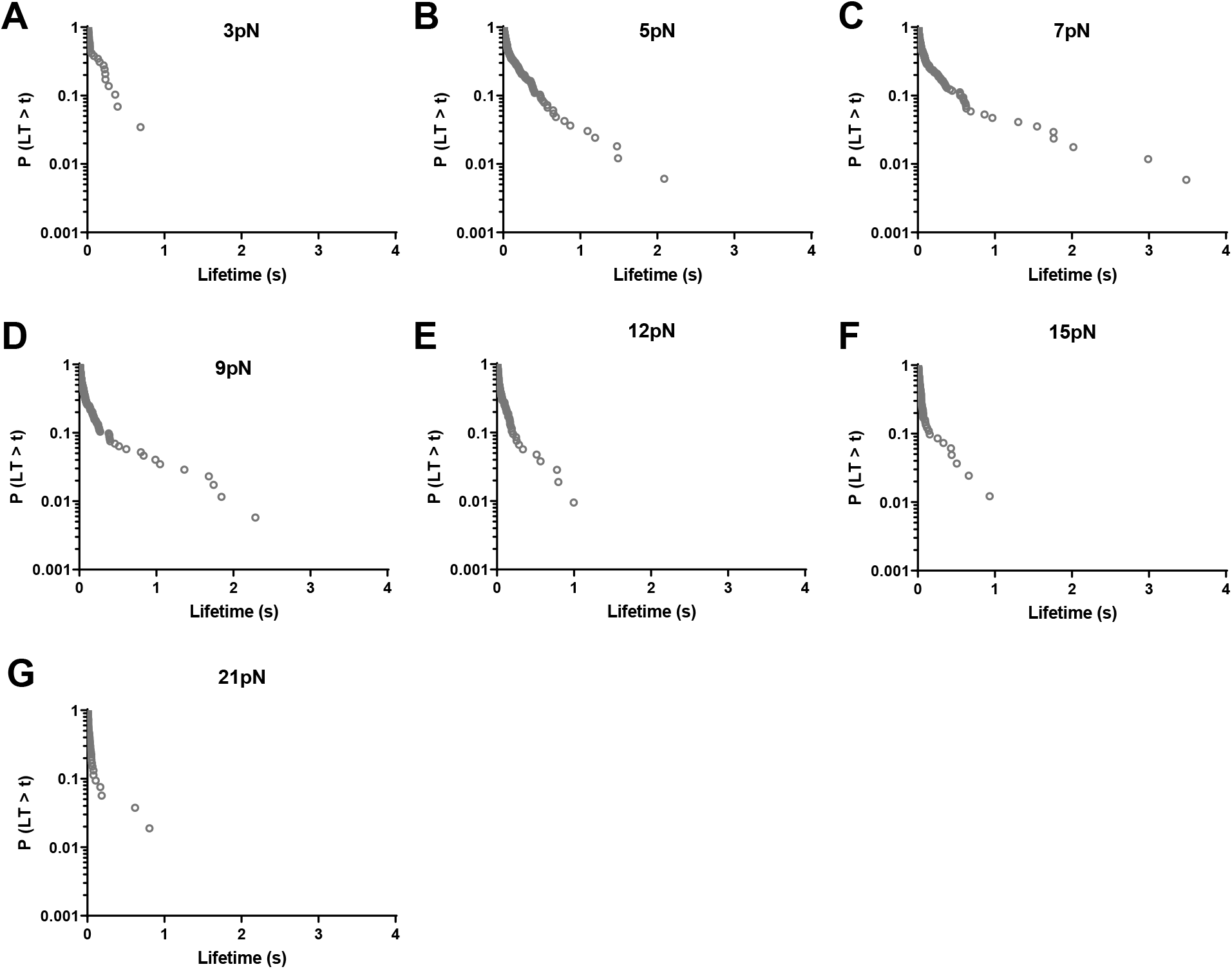
Survival frequencies (fraction of event with lifetime greater than *t*) of PD-1–PD-L2 bond lifetime for different force bins. n = 29, 165, 170, 173, 105, 82, and 53 lifetime events for (A) to (G) respectively.

